# Global composition of the bacteriophage community in honeybees

**DOI:** 10.1101/2021.09.24.461720

**Authors:** Taylor J. Busby, Craig R. Miller, Nancy A. Moran, James T. Van Leuven

## Abstract

The microbial communities in animal digestive systems are critical to host development and health. These assemblages of primarily viruses, bacteria, and fungi stimulate the immune system during development, synthesize important chemical compounds like hormones, aid in digestion, competitively exclude pathogens, etc. The bacteriophages in animal microbiomes are harder to characterize than the bacterial or fungal components of the microbiome and thus we know comparatively little about the temporal and spatial dynamics of bacteriophage communities in animal digestive systems. Recently, the bacteriophages of the honeybee gut were characterized in two European bee populations. Most of the bacteriophages described in these two reports were novel, encoded many metabolic genes in their genomes, and had a community structure that suggests coevolution with their bacterial hosts. To describe the conservation of bacteriophages in bees and begin to understand their role in the bee microbiome, we sequenced the virome of *Apis mellifera* from Austin, Texas and compared bacteriophage composition between three locations around the world. We found that the majority of bacteriophages from Austin are novel, sharing no sequence similarity to anything in public repositories. However, many bacteriophages are shared among the three bee viromes, indicating specialization of bacteriophages in the bee gut. Our study along with the two previous bee virome studies shows that the bee gut bacteriophage community is simple compared to that of many animals, consisting of several hundred types of bacteriophages that primarily infect four of the dominant bacterial phylotypes in the bee gut.

## Introduction

Honeybees (*Apis mellifera*) are the primary pollinators in agriculture, providing billions of dollars per year in pollination services. Like all animals, honeybees exist in a close-knit relationship with the microorganisms that live on and in them (1). This includes pathogens, an essential gut bacterial community, recently described bacteriophages (phages), and microorganisms living in the hive environment. These microbes are important in determining the health of honeybees and may offer a means to prevent disease, but much remains unknown about the ecological and evolutionary forces influencing the composition of the bee microbiome.

The honeybee gut microbiome is relatively simple and specialized compared to many animals. *A. mellifera* and *A. cerana*, which diverged about 5 million years ago, share five bacterial phylotypes (>97% 16S sequence identity) that are stable in relative abundance in bees across time and geographic range. These five ‘core’ phylotypes include bee-specific *Bifidobacterium*, *Snodgrassella*, *Gilliamella*, *Bombilactobacillus* spp. (previously referred to as *Lactobacillus* Firm-4), and *Lactobacillus* near *melliventris* (previously referred to as *Lactobacillus* Firm-5) (2, 3). Three additional phylotypes are often found in *A. mellifera*; *Bartonella apis*, *Frischella perrara*, and *Commensalibacter sp*., but not in *A. cerana*. At the finer taxonomic level of about 90% ANI (average nucleotide identity between whole genomes), the five core phylotypes break down into separate clusters of strains, referred to as “sequence-discrete populations’’, that can be specific to either *A. mellifera* or *A. cerana* (4). These clusters include closely related bacterial strains that likely represent distinct species, most of which lack formal nomenclature. Two species of *Gilliamella (G. apicola* and *G. apis*) have been named (5). These related species coexist in individual bee guts, and differences in their gene repertoires suggest distinct ecological niches within the bees.

The genomes of some bee gut bacteria contain an abundance of genes involved in carbohydrate metabolism and sugar transport(6, 7).These enrichments suggest that bee gut bacteria have a role in digestion and nutrient availability, compensating for processes that animals are not able to perform on their own. *Gilliamella apicola* is responsible for pectin degradation. *Bifidobacteria*, *Lactobacillus* nr. *melliventris*, and *Bombilactobacillus spp*. are responsible for the digestion of components of the outer pollen wall and coat such as flavonoids, ω-hydroxy acids, and phenolamides and for the digestion of hemicellulose and utilization of the degradation products (8). *Snodgrassella* respiration creates an anoxic environment in the hind gut, allowing for fermentation (9). Disturbances in these microbial communities, termed dysbiosis, can be detrimental (10–13). A number of studies have linked shifts in microbial composition to changes in a variety of host functions (14). Like many bacteria, CRISPR elements are present in the genomes of several honeybee gut symbionts, suggesting that bacteriophages are part of the honeybee gut microbiome (15).

Bacteriophages influence microbial communities in many ways (16). Phages facilitate nutrient cycling through host lysis (17), transfer genetic material between hosts (18, 19), encode substantial gene repertoires (20, 21), interact directly with animal immune systems (22–25), and engage in antagonistic coevolution with their bacterial hosts, impacting molecules on the surface of the cell membrane (26, 27). Although the importance of phages in microbial communities is established, much remains to be learned about phage ecology, and much of the current research is focused on the human gut and aquatic ecosystems (28). Phage populations in many other important ecosystems offer interesting study systems. Advances in genome sequencing throughput have invigorated an interest in improving our understanding of the role of microbial ecology and evolution in animal associated microbial ecosystems (22, 29–31).

Two recent papers describe phages in the honeybee gut (32, 33), providing insight into the potential influences of phages on bee gut bacteria. Bonilla-Rosso et al. (2020) sequenced phage particles from two Swiss bee colonies, analyzed total metagenomes from Swiss and Japanese bees, and isolated a handful of phages on cultured host cells. Deboutte et al. (2020) deeply sequenced phage particles from 102 hives from across Belgium. Both studies found a bee phage population that is diverse and largely unclassifiable (34). The majority of phages in the bee gut are predicted to be lytic and mainly use *Gilliamella*, *Lactobacillus* nr. *melliventris*, and *Bifidobacteria* as hosts. Most interesting is the diversity of phages observed in both studies. Whereas prophages were largely conserved across the 102 Belgian samples and in Bonilla-Rosso’s metagenomic sequencing across time, the lytic phage population was variable between samples. Many clusters of closely-related phages were also found in both studies. For example, fourteen *Bifidobacteria* phages were isolated, cultured, and sequenced by Bonilla-Rosso et al. These phages clustered into only 5 groups where the average nucleotide identity (ANI) within each group was always more than 83%. Combined, these results point towards a highly dynamic and/or rapidly evolving phage population in the honeybee gut.

Honeybees provide an ideal model for testing how phage and bacteria interact with animal hosts because of the ease of maintaining, manipulating, and reproducing large groups in controlled environments (35). However, first the temporal and geographical variation in the honeybee phage population must be understood. We sequenced phages from a colony of U.S. honeybees and compared the phage community to the two recently published studies of European honeybee colonies and found that bee phages are mostly novel, are more similar to one another than to phages from other environments, are predicted to infect only a subset of the honeybee gut bacteria, and have sequence diversity that reflects host diversity. Surprisingly, a handful of phages were highly conserved in bees from all three countries. This combination of a small set of conserved phages with a larger, highly individualized set of phages highlights a need for temporal characterization of animal-associated phages and in vitro testing of phage host range to understand animal microbiomes.

## Results

### Identifying the best viral metagenome assembly

Viral sequencing methods influence genome assembly and community characterization (36, 37). As a proof of concept, we sequenced whole-genome amplified (WGA) viral DNA extracted from Texas honeybees using Illumina MiSeq short read technology. The same sample was then sequenced using Pacbio Sequel II long read technology. Because WGA is known to introduce biases in viral community description (38, 39), we also sequenced a non-amplified sample using PacBio. Assembly of only Illumina MiSeq reads (WGA sample) resulted in 16,448 contigs with N50 3,209. The WGA sample sequenced using PacBio assembled into 9500 contigs with N50 of 14,739. The non-WGA sample was assembled into 2541 contigs with N50 of 34,304bp (Table1). These three assemblies contained overlapping sets of contigs, although the Illumina assembly was much more fragmented than either PacBio assembly, likely due to the shorter read lengths and relatively small amount of sequencing done on this library.

**Table 1.** Metagenomic assembly metrics.

|  | Input data | Read length | #Contigs | N50 | Largest contig | Mean coverage |
| --- | --- | --- | --- | --- | --- | --- |
| Illumina (WGA) | 0.7Gb | 150PE | 16,448 | 3.2kb | 0.1Mb | 1.5X |
| PacBio (WGA) | 62Gb | 4541 (N50) | 9500 | 14kb | 0.2Mb | 205X |
| PacBio (no amp) | 13Gb | 4593 (N50) | 2541 | 34kb | 0.5Mb | 77X |

Given that the non-WGA PacBio assemblies had sufficient read coverage and should be less prone to artifacts introduced by the WGA (40), we only analyzed the non-WGA assembly. Of the 2541 contigs, 412 were putatively circular molecules according to the assembler (Flye). These circular contigs ranged from 502bp to 107,828bp in length and 3X to 943X in coverage (Fig 1). In addition to the 2541 contigs, 277 plasmids were assembled by Flye. These plasmids ranged from 900bp to 33,327bp in length and from 1X to 483X in coverage. Several of these ‘plasmids’ were identified as putative phages in downstream analysis and were subsequently included in all analyses.

**Figure 1.**
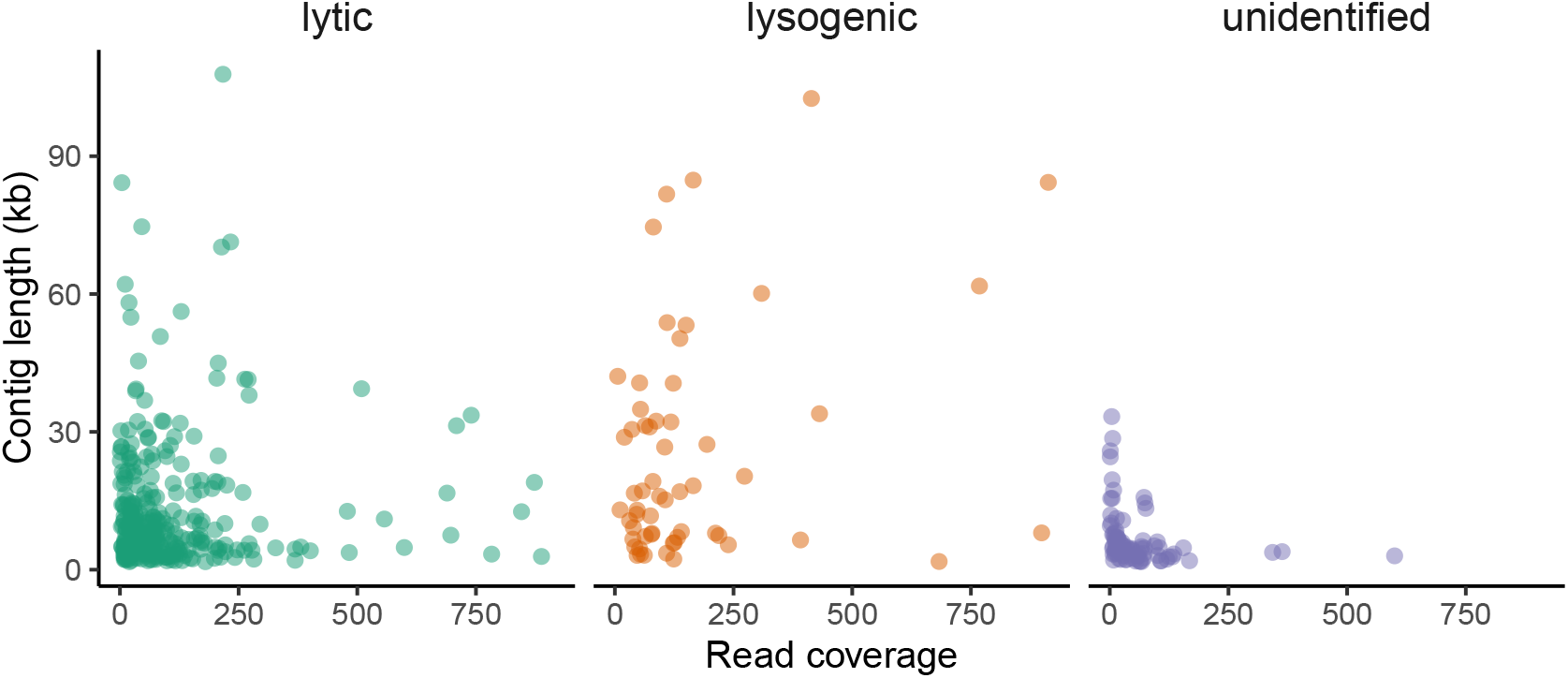
Length and abundance of contigs identified as bacteriophage. Lifestyle was determined using Vibrant.

### Identification of bacteriophage in Texas honeybees

Of the 2541 contigs in a non-WGA PacBio assembly, we identified 477 putative phage sequences (genomes) that were longer than 1000bp and were identified by at least three phage-finding programs. We found that PPRmeta identified the most phages, followed by VIBRANT, DeepVirFinder, VirSorter2, Phaster, PPRmeta, and VirfFinder, respectively (SI fig 1, SI table 1). While PPRmeta and VIBRANT identified more phages than the other programs, the phages identified by the other programs were identified by most algorithms, leading us to believe that PPRmeta and VIBRANT are the least stringent (or most sensitive) classifiers. Of the 477 putative phage, 58 were lysogenic, 325 were lytic, and 94 were unclassified by Vibrant. The length and coverages of lysogenic and lytic phages were quite variable (Fig 1) as were the assembly qualities as reported by Phaster and VIBRANT. Phaster classified 23 (18%) intact, 88 (68%) incomplete, and 18 (14%) questionable genomes. VIBRANT reported 24 (6%) high quality, 32 (8%) medium quality, and 327 (85%) low quality phage genomes (SI fig 2). These numbers seemed low considering that 214 (45%) of the contigs were identified as complete circular contigs by Flye. Combining quality and identification metrics from Phaster, VIBRANT, and Flye resulted in a list of roughly 150 phage genomes in our dataset that were high-quality (circular/complete) and had identifiable hosts or matches to phages in public repositories. The remaining 300 or so contigs (477 minus 150) had either lower quality genome completeness metrics or no host/taxonomic designatio. These contigs were still identified by at least three phage finding algorithms, so we included them in our analyses.

As was previously found by Bonilla-Rosso et al. and Deboutte et al., most phages in *A. mellifera* could not be taxonomically identified. We classified only 66 of the 477 (13%) viral contigs to the family level. These classification were based on vContact2 clustering (Fig 2) to the Viral RefSeq database and phage sequences from Bonilla-Rosso et al. and Deboutte et al. (32, 33). Bonilla-Rosso et al. and Deboutte et al. classified 24% (28/118) and 26% (73/273) of their viral clusters, respectively. The distribution of contigs into the virus families was comparable between the three studies, with the majority of phages belonging to *Siphoviridae* and *Myoviridae*, with rare observations of *Podoviridae* and *Microviridae* (Fig 3). The read coverage of individual contigs was highest for *Siphoviridae* and *Myoviridae*, further confirming their dominance in the bee microbiome. Like Deboutte et al., we did not observe any *Cystoviridae*, although one cluster of seven viral contigs contains mixed taxonomic designations that include some sequence matches to *Cystoviridae*. The hosts of these phages were predicted to be *Lactobacillus* (including the newly designated genera *Bombilactobacillus* and *Apilactobacillus*). Unlike Bonilla-Rosso et al. and Deboutte et al., we did not find any *Inoviridae*, but we did find four contigs classified as *Gokushovirinae*, all of which belong to one cluster. Two of these contigs share 99.92% identity across their ~5kb genome. Although this extreme similarity between phages was uncommon in our data, many clusters contained groups of phages sharing high sequence similarity. The vContact2 network (Fig 2) roughly illustrates the size and connectedness of these clusters. We were able to predict the hosts of 203 of the 477 phages (Fig 3) in our sample using a hierarchical scheme of searches. First, we clustered VCs from our study, Bonilla-Rosso et al. and Deboutte et al. then used host identifications from these studies. Second, we performed a blastn search against CRISPR spacer sequences from honeybee gut bacteria in Bonilla-Rosso et al. Third, we performed a blastn search against a recently compiled CRISPR sequence database (41). In both CRISPR searches, we required 100% matches between our contigs and spacer sequences. Lastly, we ran our contigs through the vHULK prediction tool (42). In only a few cases did these methods disagree, but when they did, we chose the host according to the order outlined above.

**Figure 2.**
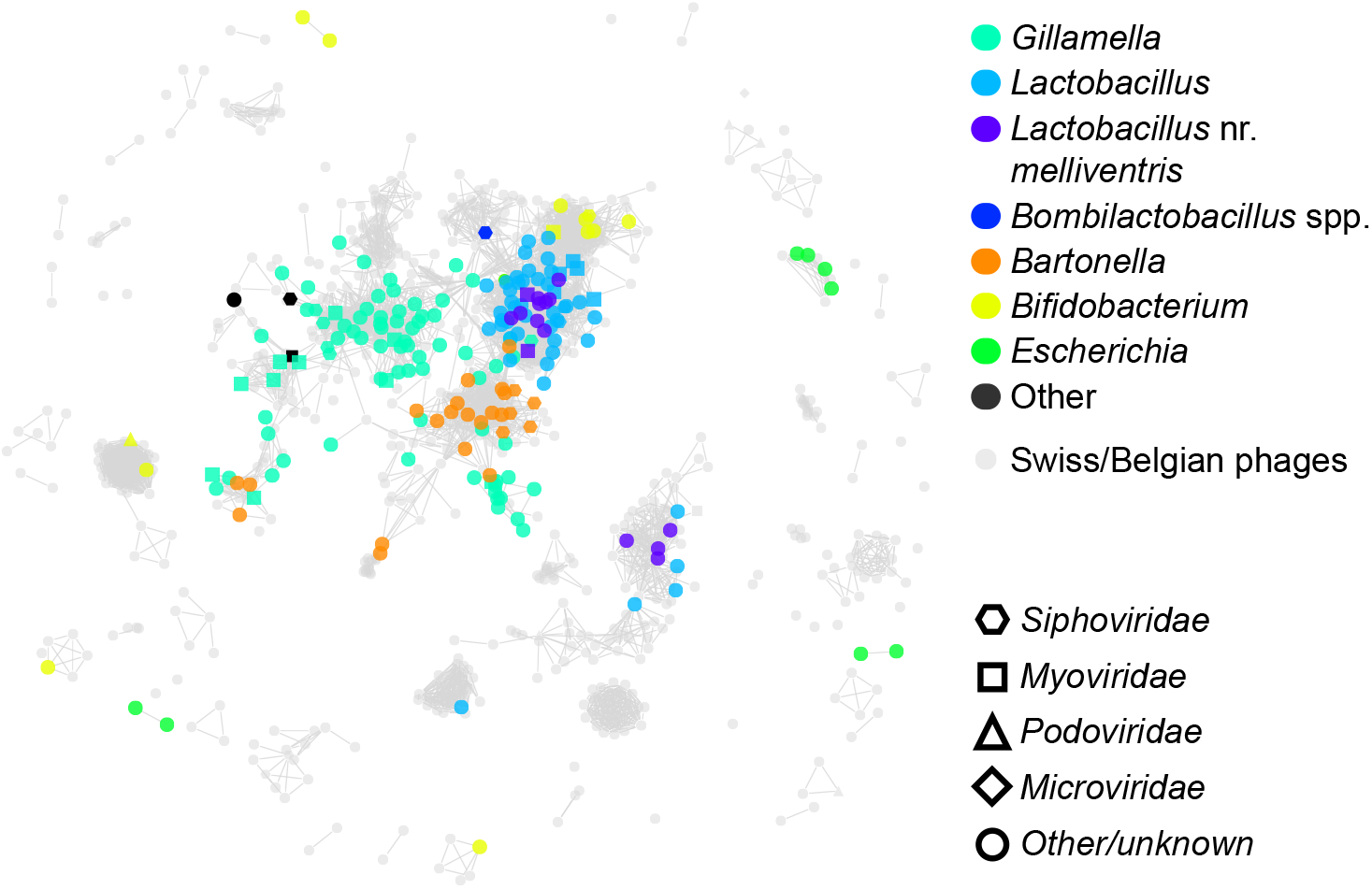
Network diagram showing clusters of Texas honeybee phages classified by host (top) and viral taxonomy (bottom). Clustering is based on similarity between protein-coding genes among viral contigs. Phages from Deboutte et al. (2020) and Bonilla-Rosso et al. (2020) are shown in grey.

**Figure 3.**
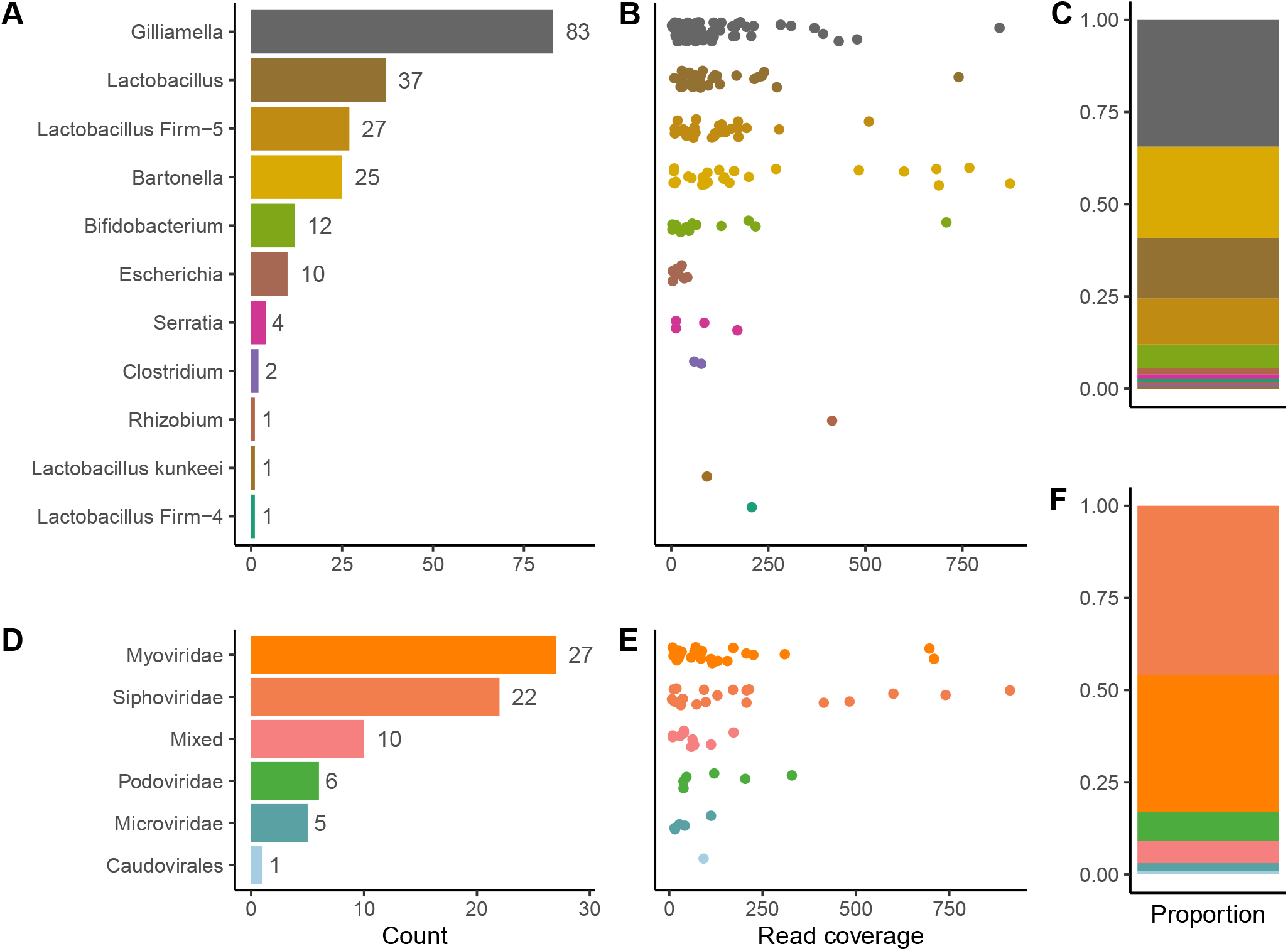
Characterization of the bee phage community. The number of phages identified is shown in panels (A) and (D). The read coverage of individual phage contigs are shown in (B) and (E). The relative abundance (read coverage) of phages by host usage and taxonomy is shown in (C) and (F).

A total of 27 contigs were predicted to infect *Lactobacillus* nr. *melliventris* and only one was predicted to infect *Bombilactobacillus*. An additional 37 contigs were classified to *Lactobacillus* broadly instead of one of the two dominant clades of *Lactobacillus-related* bacteria within honeybees (*Bombilactobacillus* and *Lactobacillus* nr. *melliventris*). Phages predicted to infect *Gilliamella* were most numerous, with 83 of the 203 phages with predicted hosts belonging to *Gilliamella*. In contrast to Bonilla-Rosso et al., and Deboutte et al., we found a number of *Microviruses* that are predicted to infect *E. coli*. We inspected these viral contigs with interest as we were worried about potential contamination from ΦX174 strains regularly used in the same lab. However, we found that there were indeed a number of *Microviruses* (sub-family *Gokushovirinae*) that were related to *Gokushovirinae* found in other bees (43). Several other *Microviruses* predicted to infect *E. coli* had unknown hosts and ~75% blastn matches to phages from a variety of environments, but not any *Microviruses* used in our lab.

### Conservation in the global honeybee phage population

Using vContact2, we clustered the 477 Texas honeybee phages with publicly available phages (Fig 2), including VCs from Deboutte et al. and Bonilla-Rosso et al. Most of the 1203 phages from the three honeybee phage metagenomic assemblies were identified as singletons or outliers by vContact2 (Fig 4, SI fig 3). These novel phages numerically dominate the bee phage community (Fig 4). The 1203 phages from honeybees clustered into 175 unambiguous clusters, 96 of which contained phages from Texas bees. A total of 184 of the 477 Texas bee phages clustered with other phages (Fig 5). Bee phages from the three honeybee viromes clustered with one another more often than with phages from viral RefSeq (Fig 4). 13 clusters contained phages from all three honeybee phage viromes. Ten of these are predicted to be lytic phages and three are predicted to be lysogenic. These phages included *Podoviruses* and *Myoviruses* of *Bifidobacteria*, *Siphoviruses* and *Caudoviruses* of *Lactobacillus* nr. *melliventris*, and *Myoviruses*, *Caudoviruses*, and *Siphoviruses* of *Gillamella*. 31 phage clusters were uniquely shared between the Belgian and Texas samples, 18 between Belgian and Swiss samples, and 8 between Swiss and Texas samples. 44 clusters from Texas honeybees did not cluster with the European bee phages but 6 of these clustered with phages in RefSeq. Many clusters that included phages from RefSeq included more than one RefSeq phage, resulting in large cluster sizes (Fig 5). The largest of these clusters contained four *Microvirids* that we identified in Texas bees but that were absent from the European samples. This cluster (VC_196) contained four *Gokushovirinae* genomes, all ranging in size from 4.5-5.5kb. Two of these differ from one another by only a few hundred nucleotides, but the others were more distantly related. All encode genes to make the major and minor capsids, internal scaffolding, replication proteins, and ssDNA synthesis proteins. While vHulk suggested an *E. coli* host, vContact2 clustered these contigs with phages identified previously in honeybees (44). Clusters containing closely related phages were common in all three datasets. The largest clusters of phages with representatives from all three bee phage viromes are mostly *Myoviruses* that infect *Gilliamella*, *Lactobacillus* nr. *melliventris*, and *Bifidobacterium* (Fig 2, 5, 6). The sequence length and synteny of phages in these clusters was quite variable. A representative set of *Myovirus* genomes illustrates the diversity observed in the vContact2 clusters (Fig 6). In the 13 clusters, the most similar phages have an average of 93% amino acid identity (AAI) across their protein coding genes (SI fig 5, SI table 5). The most dissimilar phages have 32% AAI (45% average similarity), but were mostly confined to one cluster (VC_78) of *Lactobacillus* nr. *melliventris* phages. Of the 13 clusters that contained representative genomes from all three datasets, only 5 contained Texas phages designated as being circular or being “high quality”.

**Figure 4.**
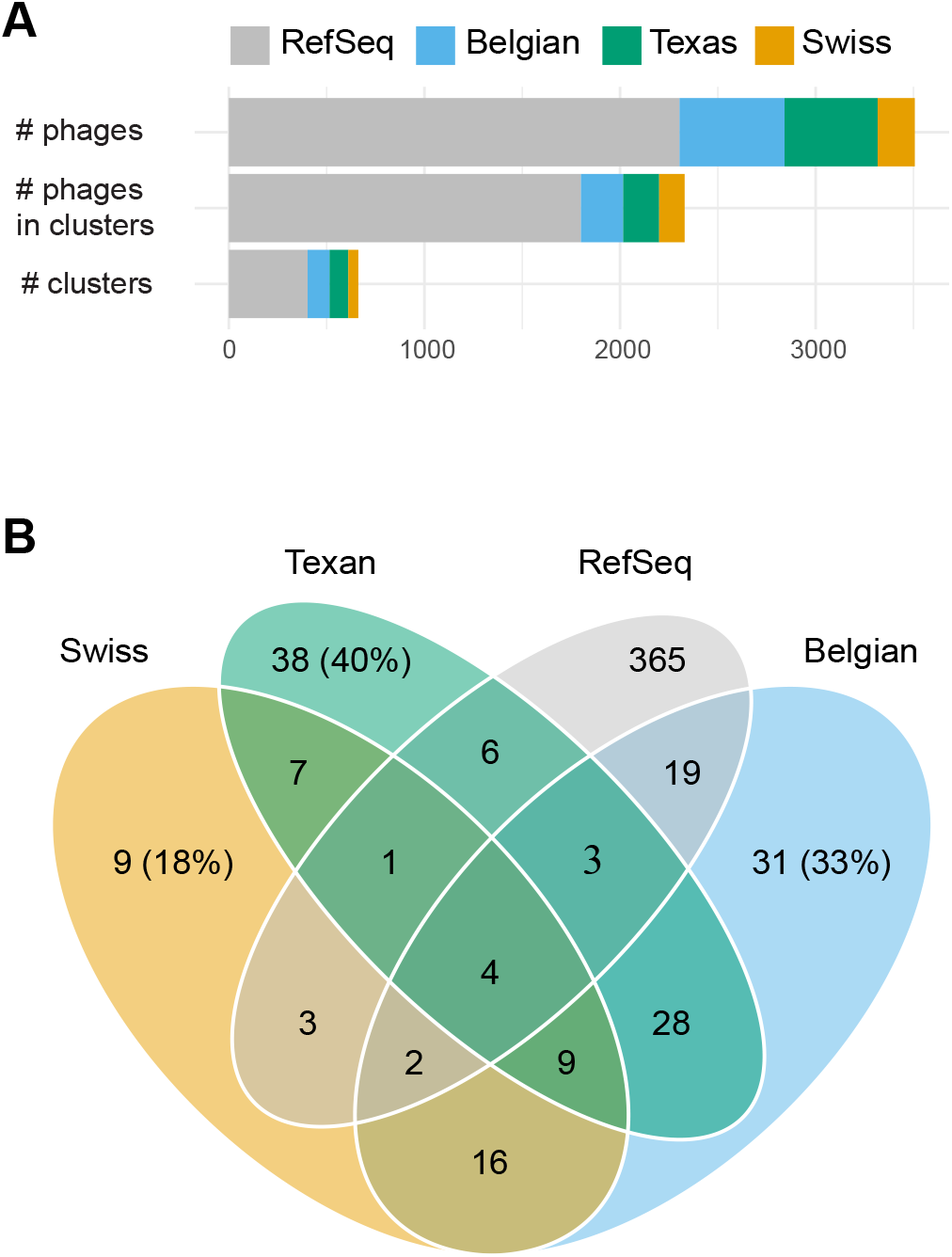
Phages from bees more readily cluster with one another than with phage sequences in reference databases. Panel (A) shows the incorporation of individual phage contigs into clusters. The composition of the final set of clusters, based on what phages are in these clusters, is shown in (B). For example, there are 9 clusters containing phages only from the Swiss data set. There are 13 (9+4) clusters with phages from all three honey bee phage viromes.

**Figure 5.**
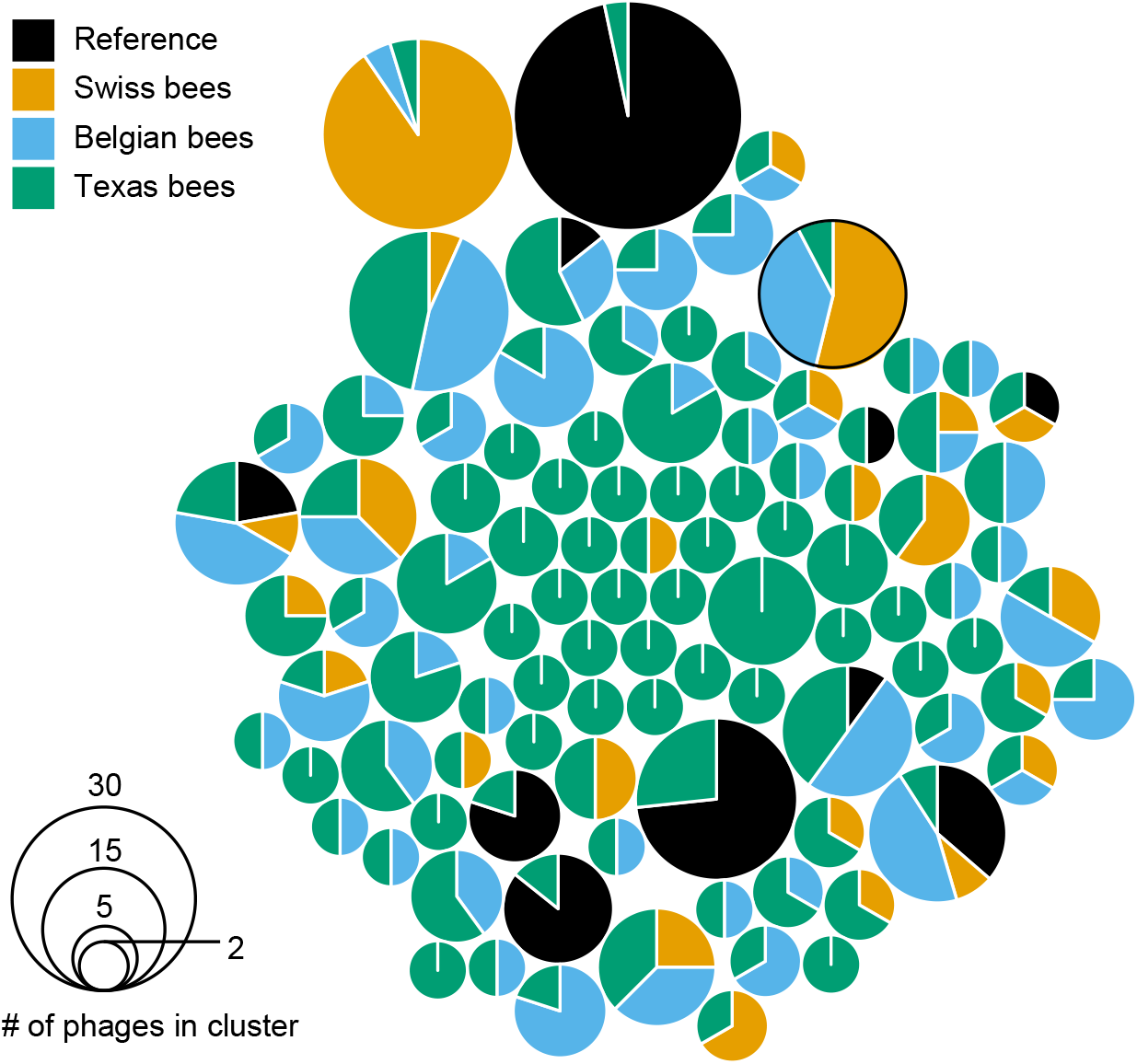
Phages from Texas honey bees mostly form small clusters of only Texas honey bee phages. A total of 96 vContact2 clusters containing 184 Texas honey bee phages contigs are shown. Circle size is proportional to the number of phage genomes in a cluster. The proportion of phage genomes from each of the four sources of phage genomes is shown. VC_106 (see Fig 6) is circled.

**Figure 6.**
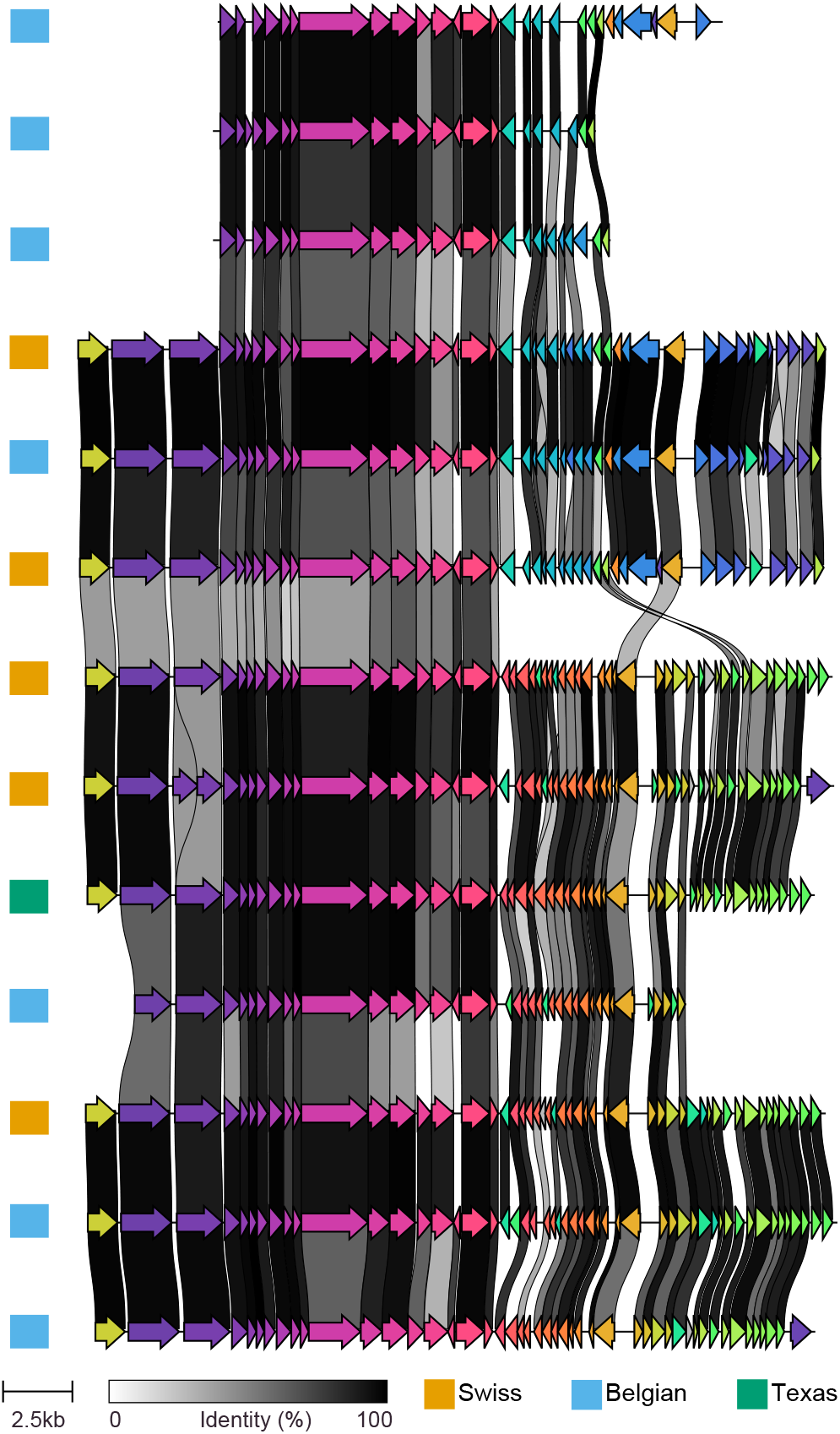
Alignment of *Bifidobacteria* infecting *Myovirus* genome from cluster VC_106. Clinker (Gilchrist et al. 2020) was used for genome ordering and visualization. A cutoff of 30% sequence identity was used for plotting connections between genes.

### Gene content of bacteriophage in Texas honeybees

A total of 15,228 protein coding genes (CDSs) were identified in the 477 phage contigs using multiPhATE2. Of these, 5983 were greater than 100 amino acids in length and 3807 had some functional annotation (i.e., not ‘hypothetical protein’ or ‘phage protein’). 48 phage contigs contained CDSs with only ‘hypothetical protein’ annotations. Of the 456 genes in these 48 phage contigs, 192 were greater than 100 amino acids in length. The number of CDSs in these contigs ranged from 3 to 31. At least some of these contigs are likely complete phage genomes containing only genes of unknown function.

The vast majority of genes with sequence matches to protein databases were phage structural proteins (capsid, tail, baseplate, portal, scaffolding). Also among the most common types of proteins were terminase, tape measure, integrase, repressor, polymerase, helicase, nuclease, DNA methyltransferase, and endolysin proteins (SI table 3). About half of the viral contigs contain at least capsid and tail/spike proteins. Other genes such as RecA, virulence factors, superinfection exclusion, and ‘plasmid proteins’ were common among the phage contigs. Metabolic genes were rare, although a handful of genes seemingly involved in queuosine, teichoic acid, and riboflavin biosynthesis were present in four viral contigs. Queuosine biosynthesis genes have been found on phage genomes and may be involved in protection from genome degradation by the host. In addition, the annotations of 372 of the 3807 CDSs were either difficult to parse (complex description) and/ or not shared among any other CDS (SI table S3). The length of these proteins was not smaller than the length of proteins with easily discernible functions, suggesting that additional functional capacity is hidden in these genes.

### Bacterial contamination of the bee virome

Contaminating bee gut bacteria were detected in the viral sequencing data at low levels. A total of 1,875 of 3,379,211 (~0.1%) of the reads were mapped to full-length bacterial small subunit ribosomal RNA (SSU-rRNA) sequences from the SIlVA database. Grouping these bacteria by genus resulted in proportional abundance of 36% *Lactobacillus* nr. *melliventris*, 17% *Bombilactobacillus*, 15% *Gilliamella*, 13% *Bartonella*, 13% *Bifidobacterium*, 2% *Snodgrassella*, 1% *Mesorhizobium* (presumably from pollen), and ~0.5% *Commensalibacter* (SI table S4). Given that phages regularly encapsulate bacterial DNA, we cannot say if these contaminating sequences are in phage capsids or simply were not digested during DNAse treatment.

## Discussion

Much remains to be learned about the ecology and evolution of bacteriophages in the microbial communities of animal digestive tracts. Even in well studied microbial communities, like in the human gut, key characteristics of the microbial ecological networks are not well understood. What phage infect what hosts? How many hosts are killed by phages? How often do temperate phages become lytic? Do phages drive bacterial diversification? Honeybees offer a promising system to study these questions. The bacterial composition of the honeybee gut is one of the most well-characterized of all animals (1, 6, 14, 35). The honeybee microbiome differs between castes (45, 46), across seasons (47), between regions (48), and changes in response to environmental conditions (10, 11). Moreover, these bacteria have beneficial impacts on their bee hosts (8) and are culturable, providing opportunity to develop microbiome-mediated treatments for disease (49).

Our comparison of the three available honeybee gut viromes provides a first look at wide-spread geographic variation in honeybee phageome, giving insight into the role of these viruses in animal microbiomes. We found that 13 phage clusters are shared among bees from Switzerland, Belgium, and the United States, suggesting that a small set of phages are widely distributed in domesticated honeybees. This set of phages is complemented by a large variable population. Interestingly, the phage community in the human digestive tract is also highly distinct among individuals with the exception of crAssPhage and perhaps a few other phages that are frequently observed in many individuals (61). crAssPhage is ubiquitous in humans and in several groups of primates (58). While the craAssphage group is diverse, some strains are highly conserved and have remained co-linear for millions of years, highlighting a paradox in phage biology: while most phages evolve quickly and are incredibly diverse, some species can be highly conserved and widely distributed.

Bacteriophages affect bacterial communities in a number of ways. Virulent (lytic) phages lyse their host bacteria, reducing their hosts’ abundance and causing the release of cellular nutrients into the environment.Temperate (lysogenic) phages integrate into their hosts’ genomes, often providing temporary benefits to their lysogens. Both virulent and temperate phages shuffle genes via horizontal gene transfer, encode auxiliary metabolic genes, and affect the evolution of their hosts. In all three honeybee viromes, virulent phages dominate the phage community in abundance and diversity. Virulent phages are often abundant in dense, productive microbial communities such as in animal (including human) digestive tracts (22, 50), moist soils (51), animal waste slurry (36), etc (30). In these types of microbial communities, phages likely play a large role in the ecology of the resident bacteria. However, we also note that strict classification of phages as virulent or temperate can be challenging given the presence of intermediate lifestyles such as pseudolysogeny, the occasional integration of phages traditionally identified as virulent, and the tendency for classifiers to identify phages as lytic (22, 52, 53). Still, the bee gut microbial community seems to be one that supports a large population of virulent phages that likely play an important role in the microbial ecology of the bee gut.

In our study, the most abundant (measured by relative read coverage) phages are not predicted to infect the most abundant bee gut bacteria, but rather *Bartonella* and *Gilliamella*. Compared to our study, Bonillo-Rosso et al. and Deboutte et al. found a higher proportion of phages infecting *Bifidobacteria* and *Lactobacillus*. None of the three studies (Texas, Belgium, Swiss) quantified the relative abundance of bacterial hosts, which could account for the differences in phage abundance. Rather host abundance is inferred from many studies on bee gut bacterial composition. Seasonal fluctuations in the absolute abundance of bacteria in bee guts can be substantial (10-100 fold) although the compositional frequencies do not change so drastically (47). The Texas and Swiss hives were sampled in January, while the Belgian hives were sampled in Autumn. Bacterial populations change from being primarily *Bartonella* and *Lactobacillus* nr. *melliventris* in the winter to *Gilliamella* and *Snodgrassella* and/or *Frischella* during foraging seasons (47). No phages predicted to infect *Commensalibacter* sp. were found in any of the three studies. Phages predicted to infect *Snodgrassella* or *Frischella* were found in Bonillo-Rosso et al. but were rare and were not found in Deboutte et al. or our study. In humans, phage and host abundances are well correlated (50). Similar correlations were observed in waste-water treatment plants (54) and seawater (55). In future studies on the bee microbiome, it will be interesting to measure bacterial and phage abundances over time to test for correlated dynamics between phage and host. Given the current data, it seems that some prevalent bacteria in the bee gut are entirely free from predation by phages.

The hosts for about 25-50% of bee phages, including one of the 13 phages present in all three bee phage studies, could not be identified, highlighting a common result in phage metagenomic studies: that phage classification can be difficult. Even in the well studied human gut, our understanding of the phage community is a moving target. Currently, crAssphages (first described in 2014) are the most abundant and diverse group of phages (56, 57). Different crAssphage genera exist across the globe and probably utilize hosts belonging to the phylum *Bacteroidetes* (58). However, finer-grain taxonomic determination of the host has not yet been resolved except in a few instances (56). Either *Microviridae* or a crAssphage-like group, *Gubaphage*, are likely the second most abundant phages (22, 25, 50), although other recently described new families are also abundant (31). Hosts for *Microviridae* and *Gubaphage* remain elusive but likely include bacteria from broad taxonomic groups. Most likely, phage abundance is dependent on many factors such as the number of available hosts, the strength of defenses employed by the host, the presence of competing phages, and the number of alternative hosts. Rapidly improving sequencing technologies (e.g., Hi-C, single cell) and host determination algorithms will help facilitate improved predictions of phage hosts and thus a better understanding of phage ecology.

Phages infecting *Gilliamella*, *Lactobacillus* nr. *melliventris*, *Bartonella*, and *Bifidobacteria* make up roughly 40%, 32%, 11%, and 6% of the unique types of assigned phages identified in our study, respectively. Many of these phages share enough sequence identity to form groups of phage clusters that are similar to one another and discrete from other bee phages. Bee gut bacteria are similarly diverse. *Gilliamella*, *Lactobacillus* nr. *Melliventris*, *Bombilactobacillus*, and *Bifidobacteria* are present in bee populations as multiple discrete clusters of related strains or species that are diverged by at least 10% across the genome (nucleotide sequence identity) (59, 60). While *Bartonella* populations also have high diversity, they do not segregate clearly into discrete clusters, based on current sampling. Individual honeybees are colonized by a small subset (sometimes just one) of the many closely-related strains present in a community. Since we sequenced a pool of 75 bees, we may have observed higher diversity of phages than are present in individual bees. Bonillo-Rosso et al. also sequenced pools of ~100 hindguts, while Deboutte et al. used smaller samples of six bees (2 bees from 3 hives each), although this required whole genome amplification. Deboutte et al. found that few phages were shared between the 102 samples they collected. The maximum number of phages shared between any two samples was 15. However, 20 phages were shared between at least 5 samples. Their sampling locations spanned the entire northern region of Belgium and were collected over 2 years. The most similar phages shared between all three bee viromes have about 93% average amino acid identity (AAI) across the genome (SI table 5). Most of the 13 clusters have phages with AAI above 90%. The most dissimilar phages (~32% AAI, 45% average similarity) were phages of *Lactobacillus* nr. *melliventris*, a diverse group of hosts. It was recently shown that closely related strains of *Lactobacillus* nr. *melliventris* coexist in the honeybee gut through niche partitioning of pollen metabolism (65). Whether or not phages specialize on these functionally divergent bacteria and if phage host range evolution affects these bacterial communities remains an interesting question. Ecological models (62) and empirical studies (63) of phage and hosts show that the stable coexistence of phages with overlapping host ranges can occur in certain conditions, specifically, when there is a fitness tradeoff between generalist and specialist strategies or in spatially-structured gut environments (64). As we learn more about phage ecology in animal guts, the bee gut microbiome provides a unique and convenient system to experimentally explore microbial ecosystems.

## Acknowledgements

We thank Holly Wichman, LuAnn Scott, Joanne Emerson, and Dan New for laboratory assistance and helpful discussions. This study was supported by the National Institutes of Health (NIH) grants P20 GM104420, NSF Idaho EPSCoR Program and by the National Science Foundation under award number OIA-1757324, startup funds awarded to CR Miller, and sequencing vouchers from The Office of Research and Economic Development at the University of Idaho. We would also like to thank the National Summer Undergraduate Research Project for matching Taylor Busby with mentor James Van Leuven, and the California Alliance For Minority Participation (LSAMP/CAMP) at University of California, Davis for providing professional support and presentation opportunities.

## Data availability

Raw sequencing reads were deposited in Genbank under accession numbers XXX-XXX. The R code for all analyzes is available in the Github repository, jtvanleuven/bee phage.

## Methods

### Preparation of sequencing libraries

Honeybees were sampled from the rooftop hives from UT Austin in January 2020. The digestive tracts of 75 *Apis mellifera* were dissected on cold PBS after euthanizing them at −20*C for 20 min. As previously described, the digestive tract is easily removed by pulling on the stinger. Remaining tissues were preserved in 95% EtOH. Approximately 7mL of PBS and dissected guts were homogenized using an ice-cold mortar and pestle. The homogenized material was centrifuged at 5000g for 5min to remove cellular debris. The supernatant was pushed through a 0.22um filter. The filters clogged after about 2mL, so 3 filters were used. The filtrate was treated with 0.25 U/uL DNAse and RNAse for 6 hours at 37*C to remove nucleic acids not protected by viral capsids. The nucleases were deactivated for 10 min using 0.5M EDTA. Nucleic acids were extracted using phenol:chloroform extraction, followed by two additional chloroform extraction steps. The volume of the aqueous layer was kept at ~5 mL by adding 10 mM Tris buffer. Ethanol precipitation using 2.5 volumes EtOH and 1/10 volume sodium acetate was used to purify nucleic acids. Contaminants were further removed using 1.5 volumes MagBio beads. The final sample was eluted in 50 uL EB. The fragment analyzer (Agilent) trace showed dilute (0.1 ng/uL) DNA of size range 100-150kb. Whole-genome amplification (GE illustra GenomiPhi V2) was performed on 1 ul purified DNA. Amplified DNA was purified with 1.5 volumes MagBio beads and sent to MiGS sequencing center for Illumina sequencing (150bp PE). Both the unamplified and WGA samples were sequenced by PacBio Sequel2.

### Sequencing

An Illumina sequencing library was generated by MiGS for the WGA DNA following Illumina’s Nextera kit protocols and was sequenced on the NextSeq 550 platform to generate 2,524,978, 150 bp PE reads.

Two PacBio libraries were made in the U of I Genomics Research Core following standard PacBio library preparation protocols. A total of 243,790 and 1,137,318 CLR reads after demultiplexing using SMRT Link (Lima) for non-amplified and WGA samples, respectively. The third library contained an *E. coli* strain closely related to ATCC 13706. We removed contigs matching this genome using bwa-mem.

### Genome assembly and analysis

Illumina reads were quality filtered and adaptor trimmed using fastp v0.20.0 (--detect_adapter_for_pe). A total of 4,983,896 filtered paired-end reads were assembled using SPADES v3.9.0 (--careful). PacBio CLR reads were assembled using FLYE with the --meta options.

FLYE-assembled contigs that were at least 2000 bp long (minimum input contig length) were analyzed in PHASTER through the web server to identify possible phage sequences. Positive hits identified as incomplete, questionable, or intact, were compared to other phage-detection software results. What the Phage was run on contigs greater than 1000 bp in length (nextflow run replikation/What_the_Phage -r v0.9.0 --cores 8 --fasta polished_1.fasta-profile local,docker). The resulting output files were combined with PHASTER results using R.

Nearly identical contigs were collapsed using cd-hit (cd-hit-i contigs_plasmids_phage.fa-c .99 -T 4 -o contigs_ plasmids_phage_cdhit99.fa). We also tested how further leniency in collapsing parameters would reduce the number of contigs and found that using a cd-hit cutoff of 95% identity reduced the 477 phage contigs down to 430 viral contigs. We choose to analyze phages at the 99% cutoff.

Phage hosts were identified using a number of tools First, vHULK v0.1 was run on contigs greater than 5kb in length. Second, blastn was used to search for Texas phage contigs in CRISPR spacer sequences from bee gut bacteria in HoneyBee-Virome-2020/pnas.2000228117. sd04.xlsx [blastn-ungapped-dust no-soft_masking false -perc_identity 100 -outfmt 6 -num_threads 4]. Thirdly, Texas phages were compared to the DASH CRISPR database using blast (blastn -db SpacersDB.fasta -query contigs_plasmids_phage.fa -ungapped -dust no -soft_masking false -perc_identity 100 -outfmt 6 -num_ threads 4). Lastly, vCONTACT2 was used to cluster phages and identify putative hosts.

A multifasta file was generated containing 477 Texas bee phages, 190 phages from Bonilla-Rosso et al. (2020), and 537 phage contigs from Deboutte et al. (2020). They were clustered using vContact2 in the CyVerse Discovery Environment (MCL clusterOne Diamond RefSeq V85 e-value=1E-4). CyVerse is supported by the National Science Foundation under Award Numbers DBI-0735191, DBI-1265383, and DBI-1743442. The vContact2 network file was visualized in Cytoscape 3.8.2 with characteristics from the combined phage identification protocol (Phaster and What the Phage). The results were visualized with (SI figure 4) and without (Fig 2) RefSeq 85 viral sequences. The clusters were arranged using the Prefuse Force Directed Open CL layout.

multiPhATE2 was used to annotate 477 putative phage genomes with the following parameters: phanotate_ calls=‘true’, prodigal_calls=‘true’, glimmer_calls=‘true’, primary_calls=‘phanotate’, blastp_identity=‘50’, blastp_hit_count=‘5’, blastp=‘true’, phmmer=‘true’, pvogs_blast=‘true’, phantome_blast=‘true’. The gene descriptions from the blastp and hmm hits were searched in R. Gene categories were roughly taken from the list of common phage genes in 10.1038/s41467-020-18236-8.

Bacterial contamination was detected by extracting PacBio reads with nucleotide similarity to small subunit RNA sequence from the SILVA database. To reduce computational requirements, only reads that were greater than 30k in length (~90% of the reads) were used (3,379,211 total reads). A condensed database was generated, clustering SSU rRNA sequences at 96% similarity following the phyloFlash protocol (66). Reads containing SSU rRNA were pulled from the metagenomic sequencing using SortMeRNA (67) using default parameters. The distribution of blast hit length and identity is shown in SI fig 4. Only full-length (1400-1700 bp) matches were analyzed for estimating bacterial contamination in the virome. MetaQUAST was used to compare assemblies from the three sequencing methods (Illumina-WGA, PacBio-WGA, and PacBio).

Gene alignments for phages in the 13 shared clusters were generated by re-annotating phage contigs using PROKKA (68) to make *.gbk files, which were then used by Clinker (69) for alignment. A shell wrapper script is included in the Github project. Analysis of the Clinker output was performed in R and is included in the Github project associated with this paper.

## Supplementary materials

Supplementary table 1. Identification of phages by nine software programs. We conservatively chose to classify a contig as a phage only if it was identified by at least three of these software programs.

Supplementary table 2. List of phage contigs and their properties including, ‘Flye.length’; contig length, ‘Flye. cov’; read coverage, ‘Flye.circ’; circular prediction, ‘IsPhage’; summary of 9 phage finding software programs, ‘Vibrant.prediction’; prediction by Vibrant, ‘Vibrant.type’; type of virus, ‘Vibrant.Quality’; genome assembly quality, ‘Phaster.COMPLETENESS’; genome assembly quality, ‘Phaster.SPECIFIC_KEYWORD’; Phaster gene matches, ‘Phaster.TOTAL_PROTEIN_NUM; number of proteins in contig, ‘Phaster.PHAGE_HIT_PROTEIN_ NUM’; number of proteins with blast hits, ‘vHulk.pred’, host prediction by vHULK, ‘VC’; cluster identification by vContact2, ‘VC.Status’; cluster classification, ‘VC. members’; contig or virus names in cluster, ‘virusTax’; taxonomic identification, ‘virusHost’; host identification, ‘crisprhit’; blast hit to bee bacteria CRISPR spacers, ‘mattInfo’; contig information from Deboutte et al. for phages that clustered with Texas phage, ‘bonillaInfo’; contig information from Bonilla-Rosso et al. for phages that clustered with Texas phage, ‘CrisprDB’; host spacer matches using the DASH CRISPR spacer database.

Supplementary table 3. Protein coding gene predictions and annotations according to multiPhATE2. Broad protein classifications are provided in the ‘parseName’ column show a simplified summary of the multiPhATE2 output.

Supplementary table 4. Summary of SSU rRNA mapping to quantify bacterial contamination of the bee virome. Columns include; SILVA reference identifier (accession number plus rRNA gene coordinates), number of matching reads, minimum, maximum, and average percent identity of the blast hit, and the taxonomic information from SILVA.

Supplementary table 5. Summary of sequence similarity between phages in the 13 clusters shared among all bee phage studies. For every pairwise comparison between contigs within each of the 13 clusters, the minimum, mean, and maximum percent identity and percent similarity values are shown for genes shared between the contigs.

