## supplementary figures for "Global composition of the bacteriophage community in honeybees"

**Supplementary figure 1.** Characterization of contigs according to what phage finding software were used for classification. Each column represents a set of contigs that were identified using the phage-finding software shown in the UpSetR plot at the bottom. Genome quality is from Vibrant. The circular designation is from the Flye genome assembler. The number of contigs identified by each set of software programs is shown as "intersection size". Most of the high- and medium-quality contigs were identified as phage by several of the software programs.

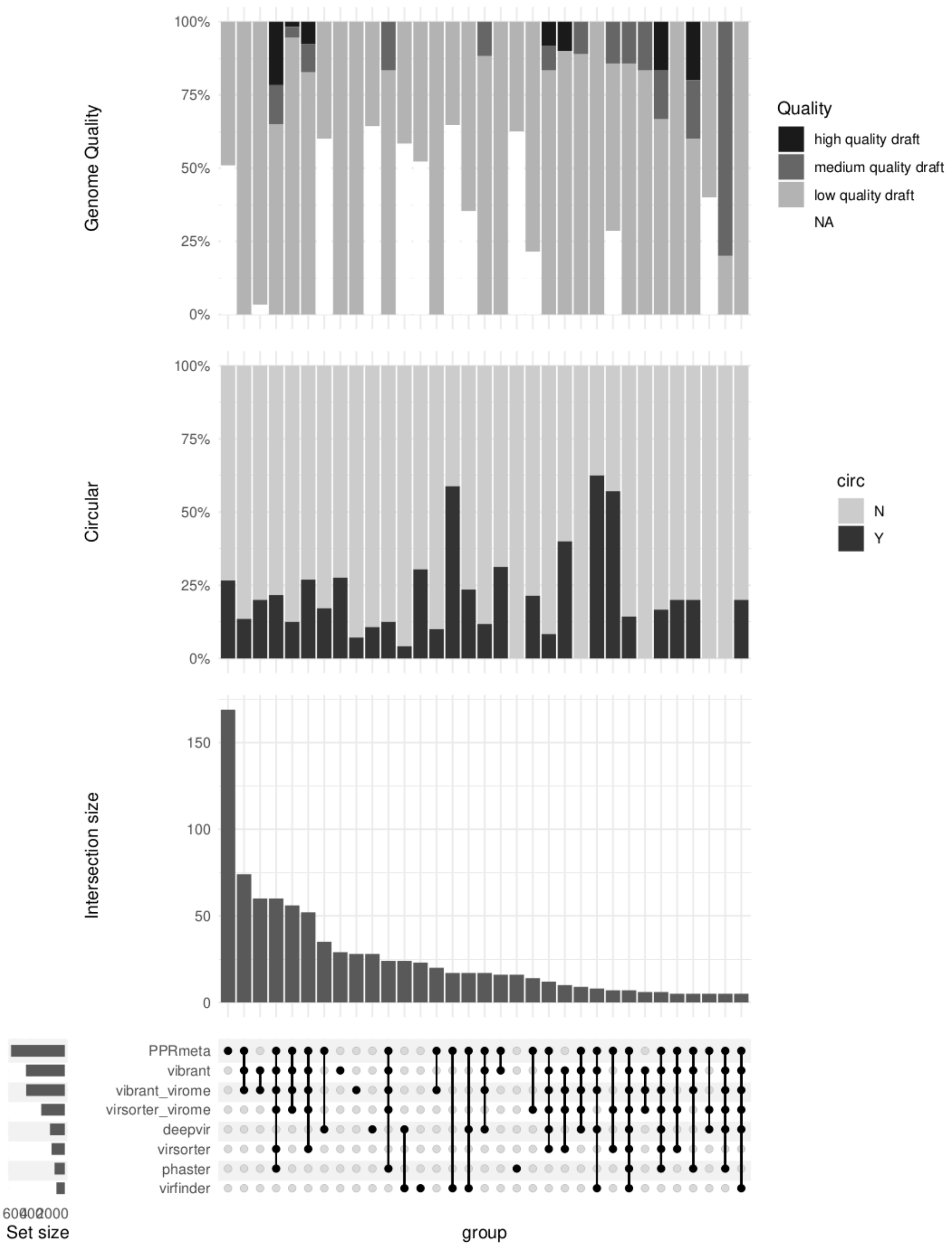

**Supplementary figure 2.** Comparison of Phaster and Vibrant genome quality metrics showing overlap in classification categories. 'NA' contigs were not identified as phages for the respective software program.

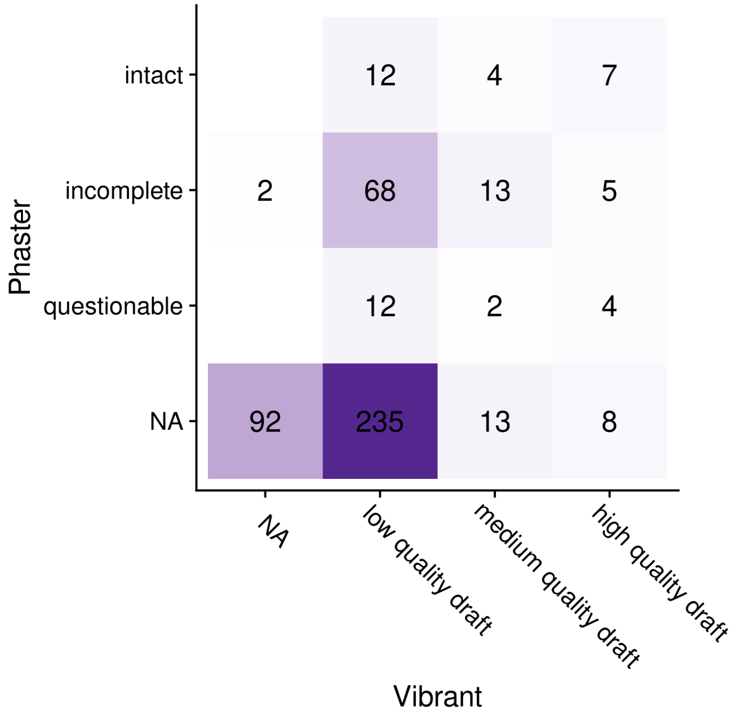

**Supplementary figure 3.** Comparison of viral contig clustering. Clustering was performed using vContact2 with viral sequences from all three bee viral metagenomes as well as Viral RefSeq. Most contigs in the 'clustered' category are clustered with other bee viral contigs and not RefSeq viruses.

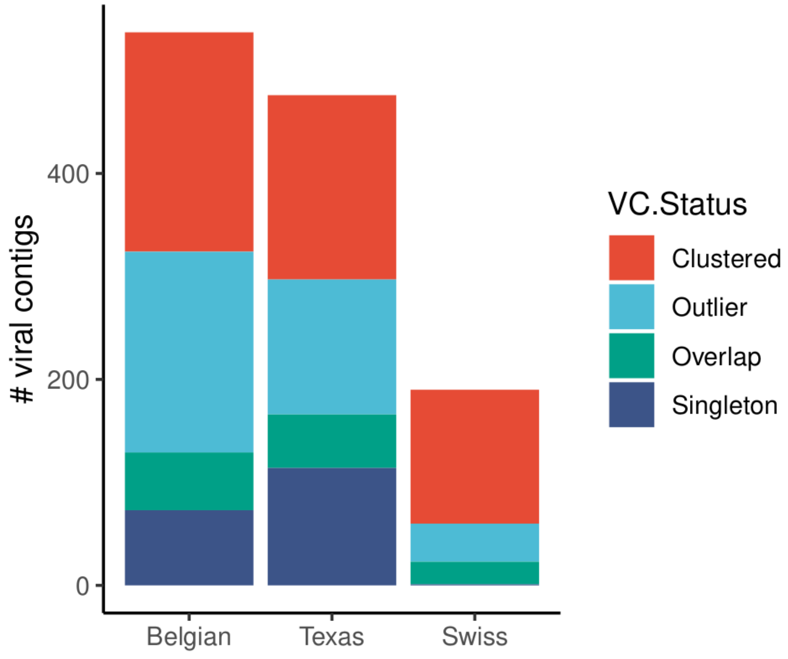

**Supplementary figure 4.** Quantification of PacBio reads containing SSU rRNA-like sequences. A total of 3,379,211 reads that were greater than 30k in length were aligned to SSU rRNAs from the SILVA database. A total of 69,666 reads had minimal e-value matches using SortMeRNA's default parameters and are plotted below. The percent identity and alignment length of the SSU rRNA blast hits are plotted. Only reads with blast hit lengths between 1400 and 1700bp were considered as bacterial matches.

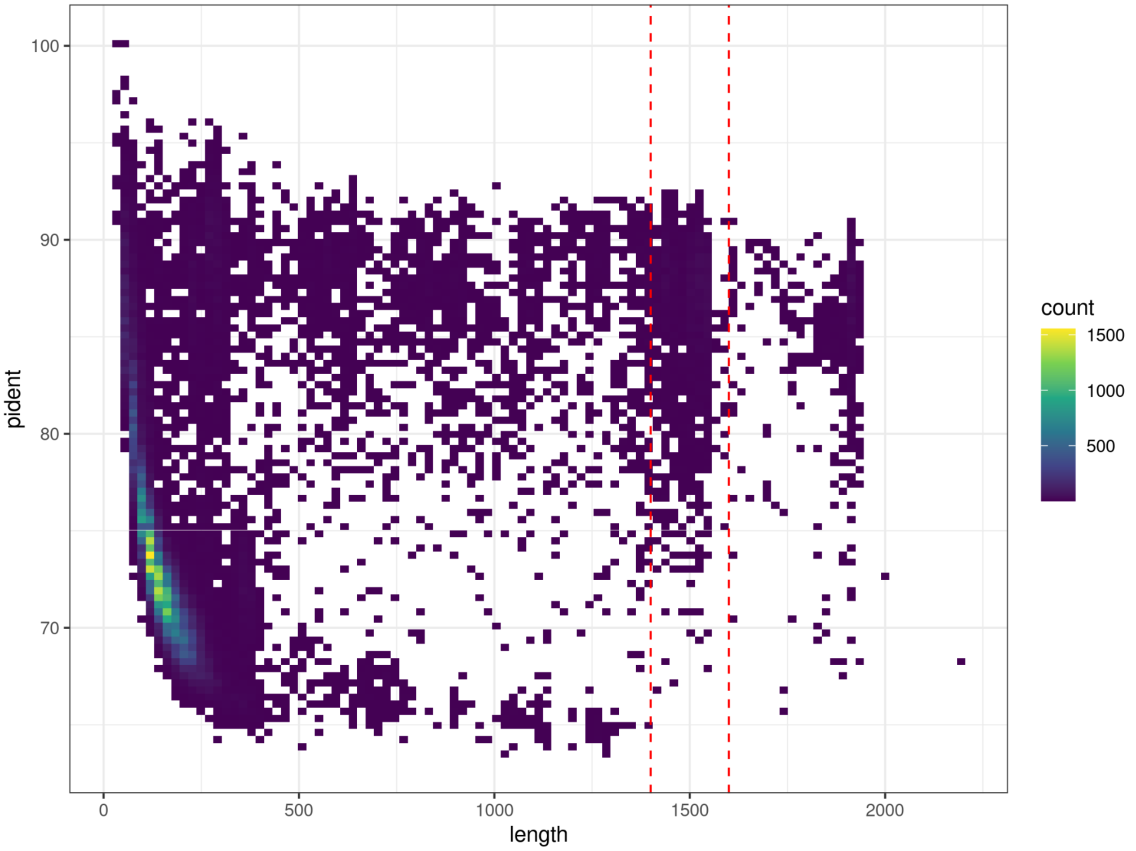

**Supplementary figure 5.** Pairwise comparisons between phages in VC_106. Histograms show the percent similarity between proteins encoded in the phage genomes. These show a large range in how similar these phages are, with some being very similar with percent similarity near 100%. Other comparisons have no proteins over ~75% similarity.

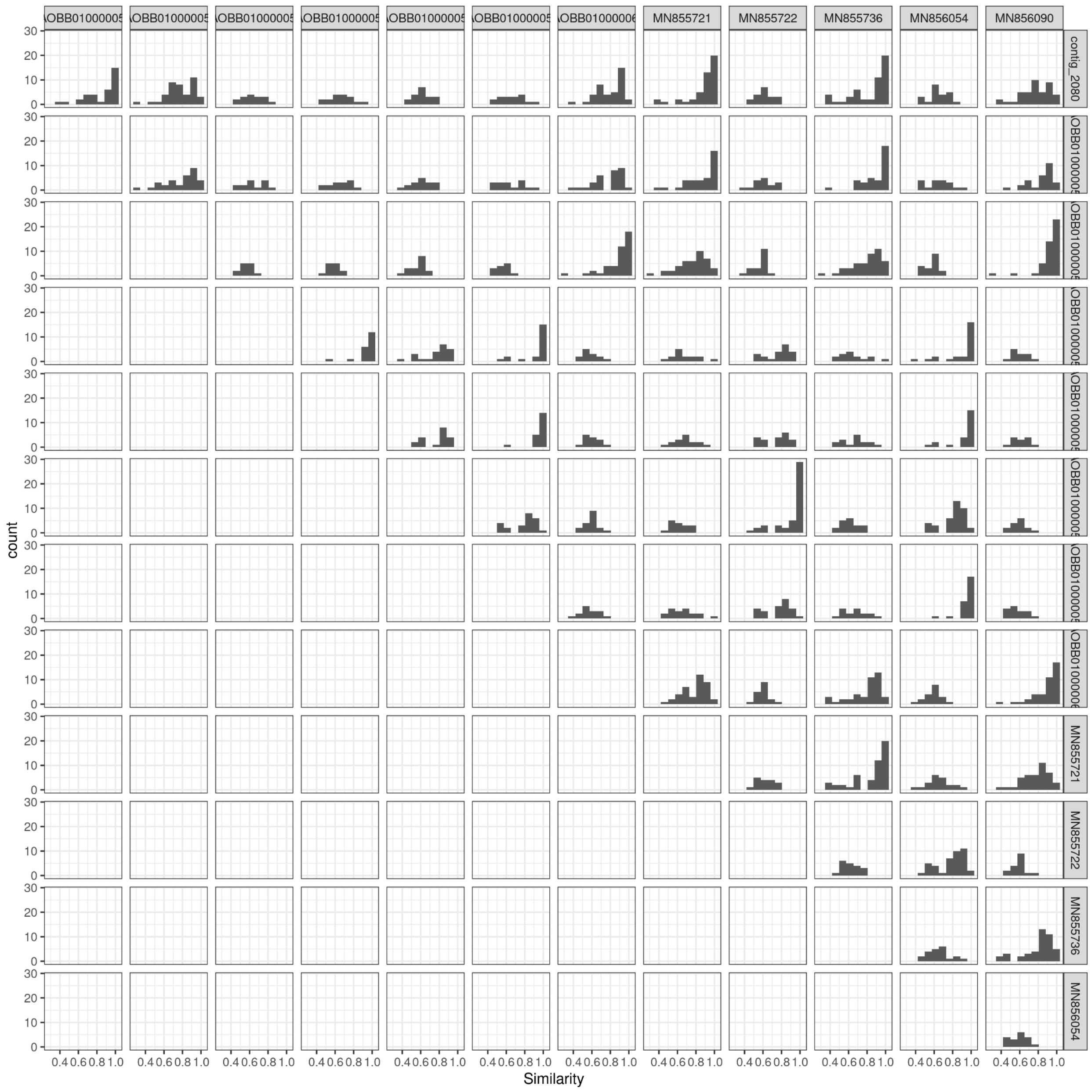
